# Dynamic integration of multisensory turn commands supports active reorienting during ongoing navigation

**DOI:** 10.1101/273805

**Authors:** Timothy A. Currier, Katherine I. Nagel

## Abstract

A longstanding goal of systems neuroscience is to quantitatively describe how the brain integrates cues from multiple modalities over time. Here we develop a closed-loop orienting paradigm in *Drosophila* to study the algorithms by which stimuli from different modalities are combined during ongoing navigation. We show that flies faced with an attractive visual and an aversive mechanosensory cue exhibit sequential responses, first turning away from the co-localized stimuli before turning back toward them. We also find that the presence of the aversive cue slows flies’ turns toward the attractive target, suggesting that conflicting unimodal turn commands are summed to produce multisensory turns of intermediate velocity. We then test a series of computational frameworks and find that integration is best described by a model in which multimodal stimuli are continuously and dynamically scaled, converted into turn commands, and then summed to produce ongoing orientation behavior.

## Introduction

A basic function of the nervous system is to combine information across sensory modalities to guide behavior. Sensory data are often synergistic, supporting similar conclusions about the state of the world, and driving similar behavioral outputs (Seilheimer, Rosenberg and Angelaki, 2013). However, conflicting cues that drive opposing behavioral responses must also be integrated by the brain (Alais and Burr, 2004; Shams, Kamitani and Shimojo, 2000). The algorithms used by the nervous system to integrate synergistic and conflicting stimuli across modalities are an active area of investigation (Fetsch et al. 2012; Drugowitsch et al. 2014, Raposo et al. 2014; Gepner et al. 2015; Burgos-Robles et al., 2017; Song et al., 2017).

The effects of multiple stimuli on behavior can be readily seen in stimulus-guided navigation, a behavior undertaken by phylogenetically diverse species, from nematodes (Lockery, 2011) and insects (Ofstadt, Zucker and Reiser, 2011; van Breugel et al., 2015) to fish (Portugues and Engert, 2009) and mammals (Buzsaki and Moser, 2013). Often, the real-world stimuli that guide navigation span many sensory modalities. When multiple stimuli that drive competing navigational responses are present simultaneously, the nervous system must resolve this conflict and select a single movement trajectory. An advantage of using navigation to study multisensory integration is that ongoing locomotion can provide a continuous read-out out of how the brain processes and combines stimuli over time (Gray, Hill and Bargmann, 2005; Gepner et al. 2015; Shulze et al., 2015).

In contrast, many studies of multisensory integration have focussed on behavioral outcomes that are localized in time. An extensive literature has examined how multisensory stimuli are integrated in order to make momentary decisions in a variety of two-alternative forced choice paradigms (Ernst and Banks, 2002; Fetsch et al. 2012; Drugowitsch et al. 2014, Raposo et al. 2014). Many studies of navigational responses to conflict have similarly focused on final outcomes, instead of examining the full time course of behavior. For example, homing desert ants and tethered flying Drosophila can adopt intermediate orientations when faced with conflicting navigational drives from vision and wind (Budick, Reiser and Dickinson, 2007; Müller and Wehner, 2007). However, it is not clear whether these intermediate orientations result from integration of desired target orientations, of turn motor commands, or of upstream sensory signals.

Another way to respond to simultaneous cues from multiple modalities is to execute competing behaviors in sequence. Sequential behaviors are commonly observed during navigation in complex multisensory environments (McMeniman et al., 2014; Lacey, Ray and Cardé, 2014; van Breugel and Dickinson, 2015). Additionally, recent studies have shown that animals can dynamically regulate their responses to constant stimuli, resulting in relatively stereotyped behavioral sequences (Seeds et al., 2014; Jovanic et al., 2016; Burgos-Robles et al., 2017). The generation of sequential responses has not been closely examined in the contexts of decision-making or navigational cue conflict.

Here we develop a novel closed-loop orienting paradigm in tethered flying *Drosophila* that allows us to examine in detail the dynamics of responses to conflicting multisensory cues. Analysis of individual flies’ trajectories revealed multiple time-varying behaviors. First, we show that flies faced with competing orientation cues exhibit sequential responses, in which simultaneous presentation of an aversive mechanosensory cue and an attractive visual cue at the same location generates first a turn away from the stimuli and then a turn back toward them. Second, we find that the presence of the aversive mechanosensory cue causes flies to turn more slowly toward the attractive visual target, suggesting that unimodal turn commands are summed to produce an intermediate turn velocity.

Based on these observations, we develop a computational model in which stimuli of different modalities are differentially and dynamically scaled, converted into turn commands, and then summed to produce ongoing orientation behavior. We show that this model can recapitulate both sequential behavior and the changes in turn kinetics that we observe experimentally. We then use the model to generate quantitative predictions of behavior in response to different stimulus intensities, and show that nonlinear “winner-take-all” algorithms do not apply in this navigational context. Finally, we spatially offset our stimuli to demonstrate that the model correctly predicts turn dynamics in response to both synergistic and competing stimuli. These results demonstrate that a system featuring summation of dynamically weighted turn commands can generate multiple, seemingly unrelated behavioral responses, and can account for previous behavioral observations in navigating flies and ants. The structure of our model argues that multimodal summation during navigation occurs on turn commands in angular velocity units, and suggests a simple and testable neural architecture for multisensory control of orientation in the insect brain.

## Results

### A continuous navigation paradigm that elicits opposing orientations to visual and mechanosensory stimuli

To generate competing navigational drives in flies, we modified a classical tethered flight paradigm (Götz, 1987) so that two stimuli of different modalities could be delivered alone or together. In this paradigm, we monitored a tethered fly’s wingbeats in real time and used the difference in wingbeat angles (a measure of intended turning behavior) to rotate a stimulus arena around the fly in closed-loop (Methods). The stimulus arena featured a high contrast vertical stripe that served as an attractive visual stimulus and an aversive wind source at the same location, which we defined as 0° (Fig. 1A). Wind and light both had rapid onset dynamics and achieved full intensity within 25 ms (Fig. 1B). Wind speed at the fly was similar across stimulus orientations (Fig. 1C).

**Figure 1.**
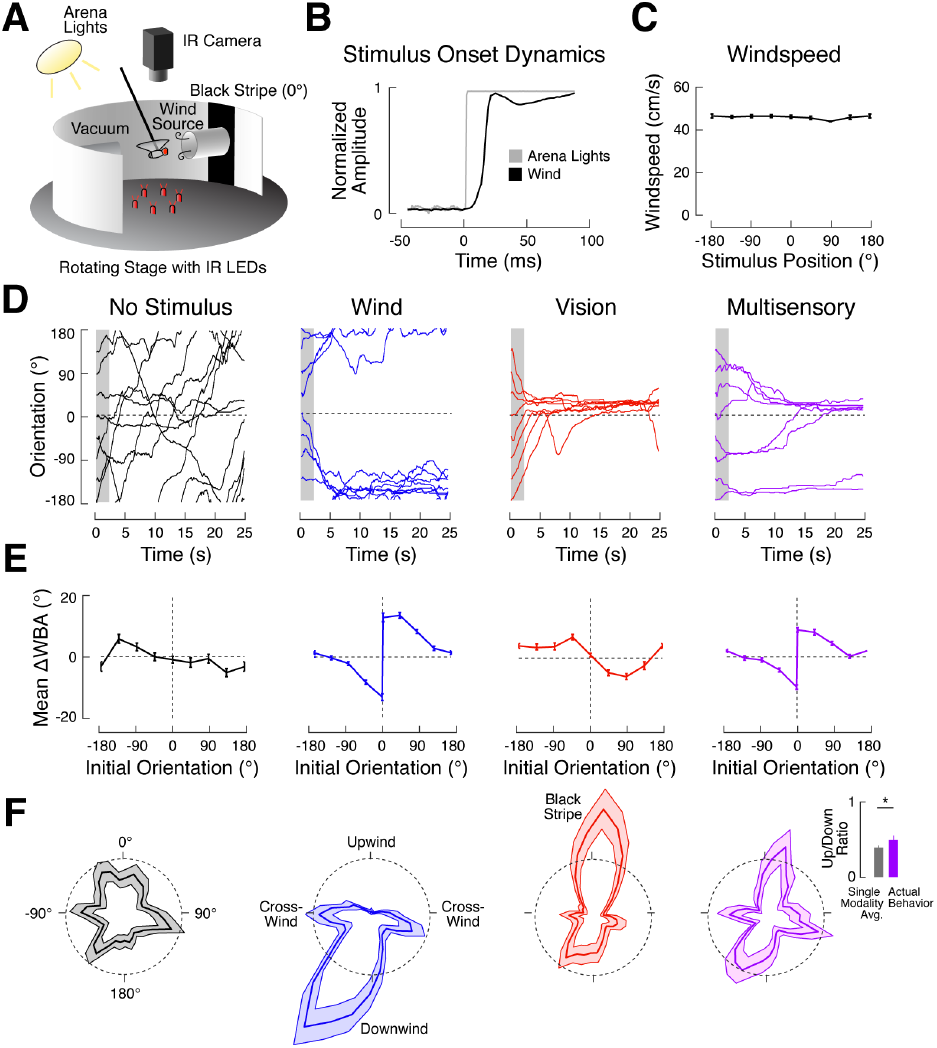
A continuous navigation paradigm that elicits opposing orientations to visual and mechanosensory stimuli. (A)Schematic of our behavioral apparatus (not to scale). A rigidly tethered fly was held between a wind and a vacuum tube. A vertical stripe was centered on the upwind tube. Attempted turning behavior was captured as the difference in wing angles (ΔWBA) with a camera and IR illumination, and was used to drive a stepper motor that rotated the arena about the fly. (B)Onset kinetics for arena lights (gray) and wind (black). Stimuli were switched on at time 0. (C) Mean windspeed +/-SEM for 8 different arena positions. Windspeed is consistent across orientations (one-way ANOVA; *p* = 0.11). (D) Example behavior of a single fly for all stimulus conditions. Each line represents a single trial. Stimulus onset at time 0. Visual stripe and wind source at 0° (dashed black lines). Data from the first 2 seconds of each trial (gray boxes) were used to calculate D-functions in (E).(E) Early trial turning behavior (D-functions) for each condition. Mean turn rate (ΔWBA) +/-SEM over the first two seconds of each trial across all flies (*N* = 120) as a function of initial orientation. (F) Late trial orientation for each condition. Polar histograms (bin width = 5°) show mean normalized orientation occupancy +/-SEM across starting orientations for the final 15 seconds of all trials. Black dashed circles represent a probability density of 0.005 per °. Orienting behavior is slightly skewed because the wind vector was not perfectly perpendicular to the edge of the arena. In the multisensory condition, the ratio of upwind to downwind orienting (inset) is higher than the mean of single modality histograms predicts (paired *t*-test, *p* < 0.05).

To investigate how flies respond to competing stimuli, we presented tethered flies (*N* = 120) with four stimulus conditions: wind only (in the dark, 45 cm/s), vision only (0.15 μW/cm^2^ arena illumination), wind and vision together, or no stimulus. Each condition was presented once from each of 8 starting orientations, randomly interleaved. After each trial, we switched off all stimuli and rotated the arena to the starting orientation for the next trial. We retained and analyzed data from flies that maintained flight through all 32 trials (45% of all flies tethered, 91% of flies that flew for at least 10 trials).

To characterize the flies’ responses to our stimuli, we first plotted orientation as a function of time for each fly (Fig. 1D). When no stimulus was present, flies had no preferred orientation, and tended to make long, slow turns. In the wind condition, flies turned away from the wind source, then maintained down- or cross-wind orientations for the remainder of the trial. Conversely, flies in the vision condition made a turn toward the stripe at stimulus onset and spent most of the trial oriented toward this stimulus. In the multisensory condition, flies also turned toward the stripe, but less reliably and more slowly than in the vision condition.

To quantitatively compare stimulus-driven turn rates and orientation preferences across conditions, we split our data into an early component where most turns occurred (first 2 s), and a late component where flies tended to maintain a stable orientation (last 15 s). To quantify early-trial behavior, we plotted flies’ mean turn rates (expressed as the mean difference in wingbeat angles) as a function of initial orientation relative to the stimuli (Fig. 1E). Historically, such plots have been called “desirability functions”, or “D-functions” (Reichardt and Poggio, 1976), a term we adopt here. The D-function for wind occupies the top-right and lower-left quadrants, indicating an aversive response to wind. For example, when the fly is oriented to the right of the wind source at +90°, it will tend to turn right (a positive-signed turn). In contrast, the D-function for vision is attractive, with data in the top-left and lower-right quadrants. In this case, a fly oriented to the right of the stripe produces a turn to the left (a negative-signed turn). In the multisensory condition, early turning resembled the wind condition, though early multisensory turns were weaker than early wind turns. Note that the wind and multisensory D-functions have their data split at 0° to emphasize that flies’ turns away from the wind source are strongest when they begin the trial pointed directly upwind, and to highlight that turns in either direction from 0° are consistent with an aversive response (Methods and Fig. S1).

To analyze late-trial behavior, we plotted orientation histograms (Fig. 1F). Flies oriented down- and cross-wind in the wind condition, and toward the black bar in the vision condition. In the multisensory condition, late-trial orientation appeared to be a combination of wind and vision behavior, with orientation up-, down-, and cross-wind. However, flies oriented upwind more than would be expected from a simple average of the vision and wind orientation histograms (Fig. 1F inset; *p* < 0.05).

Our data reveal that wind and vision in our paradigm generate opposing orienting drives. When presented together, these stimuli therefore create behavioral conflict. In response to this conflict, flies’ behavior appears to mimic the wind condition early in the trial (Fig. 1E), but is biased toward vision late in the trial (Fig. 1F). This suggests that flies may respond to conflicting stimuli by executing different behaviors in sequence.

### Evidence for sequential behavior and turn command summation in responses to conflicting stimuli

Because it is possible that different flies contribute to these early-vs. late-trial features, we next asked whether we could see evidence of sequential behavior in the trajectories of individual flies. To do this, we compared vision and multisensory trials in single flies that began at the same orientation (Fig. 2A). We focused on trials starting at 0° or 45°, as these produced little turning in the vision only condition. However, in the multisensory condition, flies first turned away from the stripe before slowly turning back toward it. The “turn away then turn back” sequential response to conflicting stimuli therefore exists in single trials.

**Figure 2.**
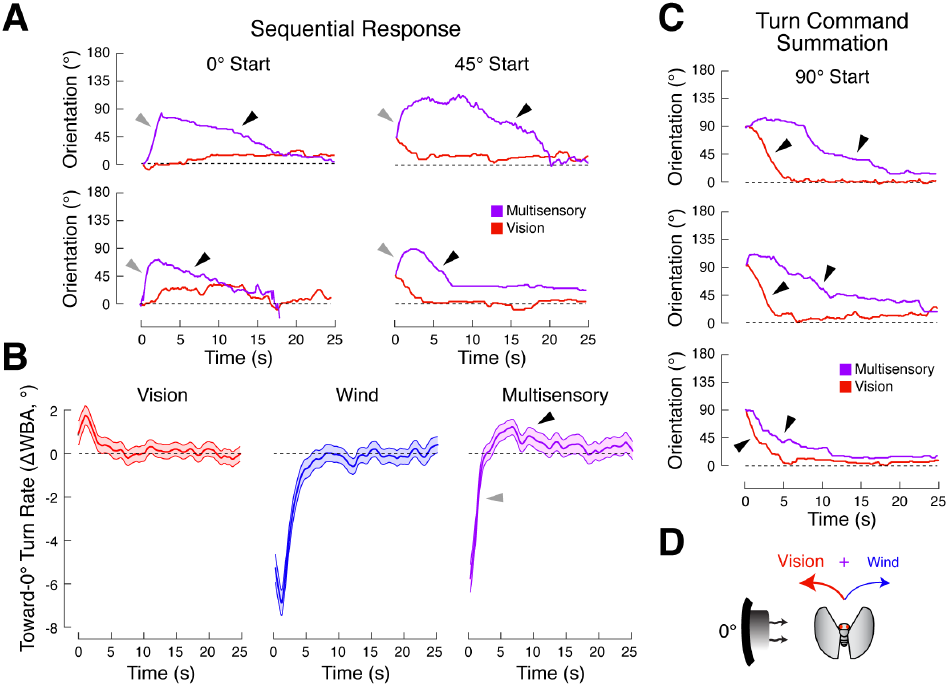
Evidence for sequential behavior and multisensory summation in single trial responses to conflicting stimuli. (A) Single trials showing evidence of sequential behavior. Each panel shows a multisensory trial (purple) overlaid on a vision trial (red) from the same fly starting at the same initial orientation. On multisensory trials, flies first turn away from 0° (gray arrows) before turning back to 0° (black arrows) later in the trial. (B) Toward-0° turn rate, expressed as ΔWBA, as a function of time in the trial for vision (red), wind (blue), and multisensory (purple) trials. Positive values indicate turns toward 0°. Traces represent mean +/-SEM for all flies at all initial orientations. Multisensory trials generate a biphasic response (gray and black arrows), characteristic of a turning sequence. (C) Single trial evidence for summation of multimodal turn commands. Multisensory trials (purple) are overlaid on vision trials (red) as in (A). Turns toward 0° are slower in the multisensory condition than in the vision condition (compare slopes at black arrows), suggesting that a negative wind-based turn command has been added to a positive vision-based turn command to produce a slower overall turn. (D) Schematic of multimodal turn command summation (not to scale). A large visual turn command (red) drives the fly left while a smaller wind-based command (blue) drives the fly right. On multisensory trials, these commands are summed to produce a reduced-magnitude turn in the direction of the larger command.

To assess the generality of this turning sequence across flies and starting orientations, we plotted the toward-0° turn rate (Fig. 2B). Here, all turns toward 0° are positive, and all turns away from 0° are negative. In the vision condition, the toward-0° turn rate shows a small positive peak near stimulus onset (Fig. 2B, left). Conversely, the wind condition data shows a large negative peak (Fig. 2B, center). In the multisensory case (Fig. 2B, right), the toward-0° turn rate is biphasic, with an early negative component and a later positive piece, consistent with an early turn away from 0° and a later turn back toward 0°. This analysis demonstrates that the sequential response is a general phenomenon across starting orientations.

Examination of individual trajectories revealed a second phenomenon in flies’ responses to multisensory competition: a change in turn rate on multisensory trials. In many cases, flies turned toward the stripe in both the vision and multisensory conditions, but the multisensory turn rate was slower than the visual turn rate. The change in turn rate was particularly evident in trials starting at 90° (Fig. 2C), but was also visible in trials starting at other orientations (Fig. 2A). These observations suggest that turn commands driven by the two modalities may be summed (Fig. 2D). For example, a vision command that drives the fly toward the stripe may be summed with an oppositely signed wind command, producing a slower multisensory turn.

### Multisensory summation and sequential responses are consistently observed across flies and require antennal mechanosensation

We next asked if sequential responses and multisensory summation were consistent features of flies’ behavior. We presented 11 additional flies with wind (25 cm/s), vision (15 μW/cm^2^), and multisensory trials starting at either 90° or 0° five times each. We chose 90° because it most clearly elicited evidence of summation (e.g., Fig. 2C), while 0° elicited the clearest evidence of sequential behavior (e.g. Fig. 2A).

As in previous experiments, we found that flies’ turns from 90° toward the stripe were slower in the multisensory condition than in the vision condition (Fig. 3A-B). To quantify this observation, we compared the latency and speed of turns toward the stripe on vision and multisensory trials. We found that the latency to cross 45° (halfway through turns from 90° to 0°) was longer on multisensory trials (Fig. 3C, left; *p* < 0.01), supporting the hypothesis that these turns are slower. Mean turn rates over the second surrounding 45° crossings were also slower on multisensory trials (Fig. 3C, right; *p* < 0.05).

**Figure 3.**
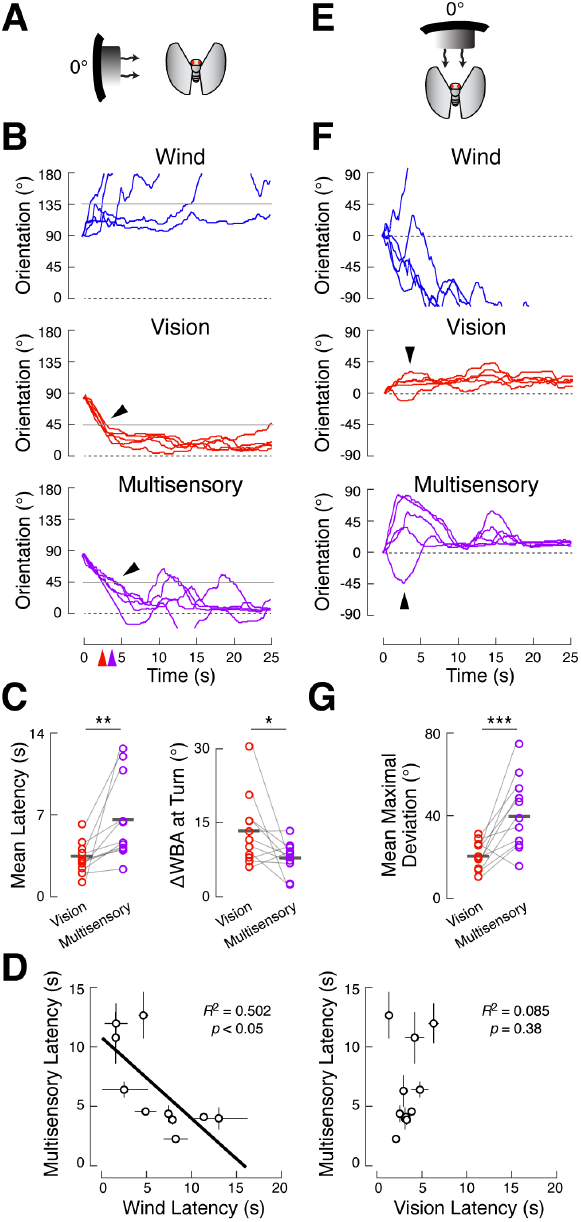
Multisensory summation and sequential responses are consistently observed across flies. (A) Stimuli starting at 90° to the fly elicitedevidence of turn command summation. Eleven flies received 5 presentations of wind (25 cm/s), vision (15 μW/cm^2^), or both. (B) Example behavior of a single fly. Orientation axis is truncated and time axis is expanded for clarity. Turns toward 0° in the multisensory condition are slower than those in the vision condition (black arrows). Gray lines indicate the midpoints of stimulus-guided turns (45° or 135°) used to calculate turn latency and speed in (C). Colored arrows on the time axis show the mean latency to cross 45° for vision (red) and multisensory (purple) trials in this fly. (C) Left: mean latency to cross 45° across 5 stimulus presentations for all flies in the vision (red) and multisensory (purple) conditions. Gray bars: means across flies. Multisensory turns occurred later than vision turns (paired *t*-test, *p* < 0.01). Right: mean turn rate, as ΔWBA, over a 1 s window centered on 45° crossings for each fly. Multisensory turns are slower than vision turns (paired *t*-test, *p* < 0.05). (D) Left: individual differences in wind sensitivity (x-axis) predict multisensory turn latency (y-axis). Each circle represents the mean latency to cross 135° in the wind condition versus the mean latency to cross 45° in the multisensory condition (as in (C)) for each fly. Bars represent SEM for each measurement. Greater wind sensitivity (small wind latency) was correlated with slower multisensory turns (*R*^2^ = 0.50, *p* < 0.05; 10 of 11 flies reliably turned through 135° in the wind condition and were included in the analysis). Black line: best linear fit to data. Right: individual differences in visual turn latency (x-axis) do not predict multisensory turn latency (y-axis). Each circle represents the mean latency to cross 45° in the vision condition versus the mean latency to cross 45° in the multisensory condition. No correlation was observed between these measures (*R*^2^ = 0.09, *p* = 0.38; 10 of 11 flies showed reliable turns through 45° in the vision condition and were included in the analysis). (E) Stimuli starting at 0° to the fly elicited evidence of sequential behavior. (F)Example behavior of a single fly showing larger deviations from 0° in the multisensory condition than in the vision condition (black arrows). (G) Mean maximal deviation from 0° for all flies in the vision and multisensory conditions. Maximal deviations are the largest absolute orientation attained over the first 10 seconds of each trial. Flies turned farther from 0° in the multisensory condition than in the vision condition (paired *t*-test, *p* < 0.001).

If turns in the multisensory condition reflect the sum of oppositely signed vision and wind turn commands, then flies with larger wind command magnitudes should show the slowest multisensory turns. In separate experiments, we found that flies differed in their sensitivity to wind (Fig. S2). We therefore asked whether single-fly wind sensitivity predicted the extent of visual turn slowing on multisensory trials. We plotted each fly’s multisensory turn latency as a function of that same fly’s wind turn latency (Fig. 3D), calculated as the average time for each fly to cross 135° (halfway through turns from 90° to 180°) on wind trials. We found a negative correlation between these measures (*R*^2^ = 0.50, *p* < 0.05), indicating that the flies most sensitive to wind, with the largest wind turn commands, take the longest to turn toward the visual stimulus in the multisensory condition. Surprisingly, multisensory turn latency was not significantly correlated with latency in the vision condition (*R*^2^ = 0.09, *p* = 0.38), likely due to the small variance in vision latency. This result suggests that differences in wind sensitivity are the larger determinants of multisensory turn rate. Together, our results provide quantitative evidence for summation of multisensory turn commands, where the magnitude of the visual command is reduced by the addition of an oppositely-signed wind command, and the magnitude of that wind command predicts the extent of turn slowing.

We next evaluated whether single flies consistently produce a turning sequence on multisensory trials by examining trials starting at 0° (Fig. 3E). In these conditions, flies first turned away from the stripe before returning to it, indicating a sequential response (Fig. 3F). To assess the degree to which each fly performed this sequence, we computed the maximal deviation from 0° over the first ten seconds of each trial (Fig. 3G), a more robust measure than the toward-0° turn rate for trials starting at 0°. On vision trials, flies remained oriented toward the black stripe and therefore had small maximal deviations. The same flies turned farther from 0° on multisensory trials (*p* < 0.001), consistent with the idea that sequential responses are common across flies. Collectively, our results indicate that multimodal summation and sequential responses are consistent features of flies’ responses to navigational conflict.

The behaviors described above might arise from neural computations, or they might arise from mechanical forces acting on the fly’s body or wings. For example, the force of the wind might make it hard for flies to make rapid turns toward the stripe on multisensory trials, or might force the fly initially off course, resulting in a turning sequence. To discriminate between these possibilities, we stabilized the antennae of 12 flies (Fig. 4A) and repeated the experiment described in Figure 3. Antennal movements are required for transduction of wind direction information (Budick, Reiser and Dickinson, 2007; Yorozu et al. 2009) and antennal stabilization abolished downwind turning and orienting in our paradigm (Fig. S3). If sequential responses and multimodal summation rely on neural computations, antenna-stabilized flies should not show these behaviors. Alternatively, if they arise from mechanical forces acting on the body, these behaviors should still be present in antenna-stabilized flies.

**Figure 4.**
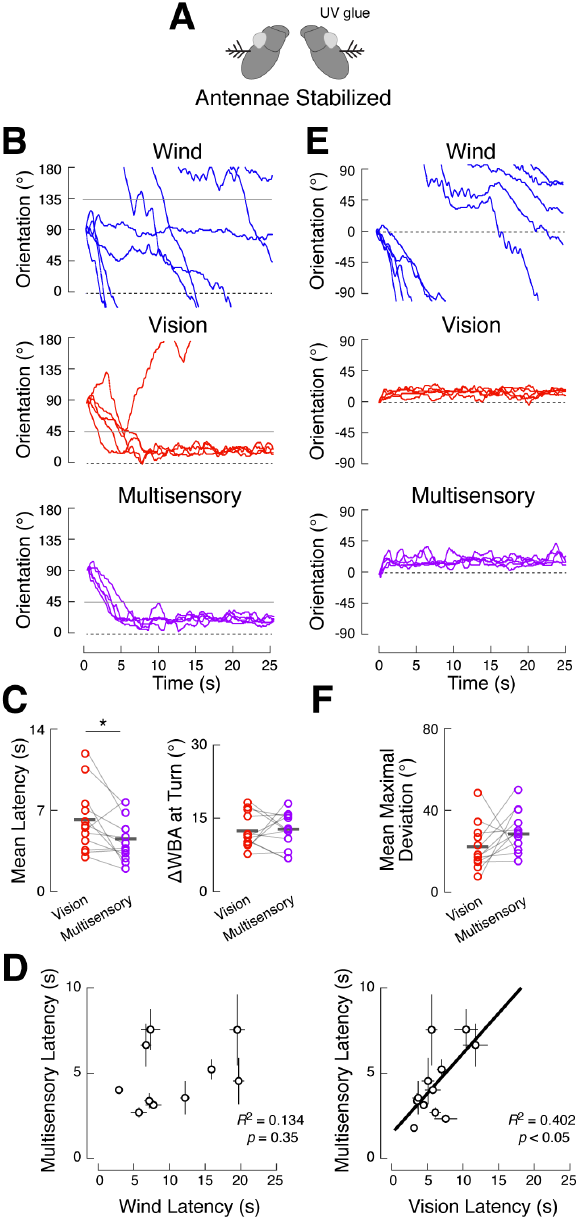
Multisensory summation and sequential responses require antennal mechanosensation. (A) Schematic of antenna stabilization procedure, using a drop of UV-cured glue at the junction of the second and third antennal segments (see Fig. S3) (B) Example behavior of a single antenna-stabilized fly in response to stimuli starting at 90°, plotted as in Fig. 3B. Turns toward the stripe are similarly fast in the vision and multisensory conditions. (C) Latency and turn rate for vision and multisensory trials in 12 antenna-stabilized flies, plotted as in Fig. 3C. Left: in antenna stabilized flies, multisensory turn latency is faster (paired *t*-test, *p* < 0.05) rather than slower, as seen with intact antennae. Right: mean turn rate of antenna-stabilized flies in vision and multisensory conditions are not significantly different (paired *t*-test, *p* = 0.93). (D) In antenna-stabilized flies, individual differences in wind sensitivity (left) are not correlated with multisensory turn latency (*R*^2^ = 0.13, *p* = 0.35), but vision latency is correlated with multisensory latency (*R*^2^ = 0.40, *p* < 0.05), consistent with a loss of the wind turn command. Plots as in Fig. 3D. (E) Example behavior of a single antenna-stabilized fly in response to stimuli starting at 0°. The fly shows no turning sequence in the multisensory condition. (F) Mean maximal deviation from 0° in the vision and multisensory conditions for all antenna-stabilized flies, plotted as in Fig. 3G. No significant difference is observed between conditions (paired *t*-test, *p* = 0.11).

Antennal stabilization abolished the slowing of visually-guided turns that we observed in the multisensory behavior of control flies (Fig. 4B). In trials starting at 90°, turn rate was not significantly different between the vision and multisensory conditions (Fig. 4C, right; *p* = 0.93). In control flies, we found that wind sensitivity predicted multisensory turn latency, but this correlation was not present in antenna-stabilized flies (Fig. 4D; *R*^2^ = 0.13, *p* = 0.35). Instead, visual sensitivity predicted multisensory latency (*R*^2^ = 0.40, *p* < 0.05), consistent with antenna-stabilized flies lacking a wind turn command, leaving only the vision command to influence multisensory turn rate. Curiously, while control flies took longer to execute visually-guided turns in the multisensory condition compared to the vision condition (Fig. 3C, left), antenna-stabilized flies executed their turns more *quickly* in the multisensory condition (Fig. 4C, left; *p* < 0.05) perhaps reflecting the input of non-antennal wind-sensors (Fig. S3).

Stabilizing the antennae also abolished sequential responses to competing stimuli (Fig. 4E). On average, antenna-stabilized flies turned no farther from the stripe on multisensory trials compared to vision trials (Fig. 4F *p* = 0.11). The lack of evidence for both multimodal summation and sequential behavior in antenna-stabilized flies argues that these behaviors rely on neural computations and on the presence of multiple stimulus-driven turn commands within the brain.

### A behavioral model that captures multimodal summation and sequential responses suggests a neural “order of operations”

Our experimental data indicate that flies respond to conflict between multisensory navigational drives with either sequential turning or summation of single modality turn commands. However, which response flies make appears to depend on the initial orientation of the competing stimuli. We wondered if a single computational framework could account for both behavioral observations and predict behavior across the full range of stimulus positions. We therefore developed a simple model of multisensory integration.

In this model, the angular velocity at each point in time is specified by a sum of single modality D-functions - the turn command produced by each stimulus given its orientation relative to the fly. This configuration produces multimodal summation, as multisensory angular velocity is the sum of the turn commands produced by wind and vision. To account for sequential responses, we reasoned that some aspect of the model must change over time. Because flies make downwind turns early in multisensory trials, the relative weight flies give to wind is likely to be high early in the trial. Over time, however, flies cease their downwind turns and eventually execute upwind turns toward the stripe, suggesting that either the relative weight of wind decays over time, or the relative weight of vision grows over time, or both. Thus, we created time-varying terms that scale the magnitude of each D-function over time: the wind weight starts high and decays over time, while the visual weight starts low and grows over time. Static scaling terms, α, represent stimulus intensity for each modality. Multisensory behavior therefore reflects a dynamically-weighted sum of unimodal turn commands (Fig. 5A).

**Figure 5.**
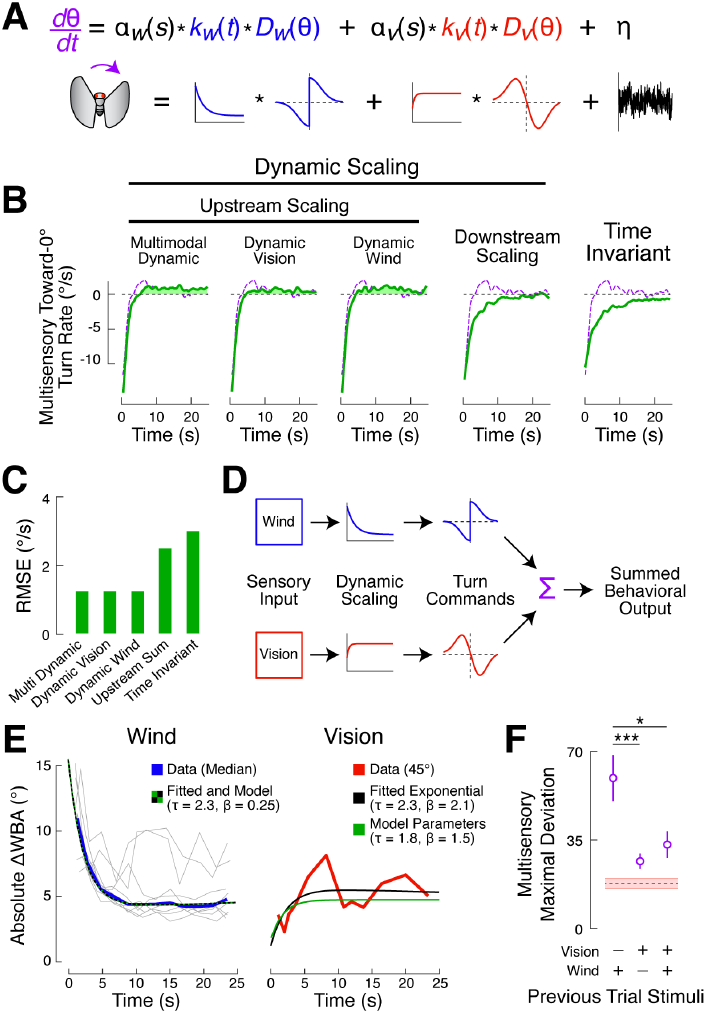
A behavioral model that captures multimodal summation and sequential behavior suggests a neural order of operations. (A) Formulation and schematic of the model. Instantaneous multisensory turn rate (*d*θ/*dt*) is given by a sum of single modality D-functions, *D*_*w*_ (blue) and *D*_*v*_ (red), weighted by static stimulus intensity variables (α), and dynamic scaling terms (*k*) that generate exponential growth or decay. A noise term (α) matched to the spectral properties of our data was added to generate simulated behavioral traces in Figs. S5, 6, and 7. Model details in Methods, Table 1, and Figure S4. (B) Measured (purple dashed lines, replotted from Fig. 2B) and predicted (green) multisensory toward-0° turn rate for the model shown in (A) and for various simpler models. A biphasic toward-0° turn rate indicates sequential turning and requires at least one modality to change over time relative to the other (achieved only by “upstream scaling” models). The positive portions of the model traces are filled in for clarity. (C) Quantification of model fits. Root mean-squared error between each model’s best-fit simulation and the experimental toward-0° turn rate in the multisensory condition. Upstream scaling models fit the data better than simpler versions. (D)Schematic of the neural “order of operations” suggested by our model: sensory inputs are independently and dynamically scaled, converted to turn commands, and summed. Scaling and turn command generation both occur prior to summation. (E) Left: absolute turn rate as a function of time for 120 flies (from Figs. 1 and 2) passing through a range of orientations (30°, 45°, 60°, 75°, 90°, 105°, 120°, 135°, and 150°) in the wind condition (gray) is overlaid with the median of these curves (blue) and a line that represents both the best exponential fit to the median and the wind scaling function, k_w_, from the multimodal dynamic model (black and green). The parameters returned when fitting the blue curve were identical to the best-fit parameters in the multimodal dynamic model. Right: absolute turn rate as a function of time for the same flies passing through 45° on visual trials that were preceded by lights-off (wind) trials (red) is overlaid with the best exponential fit to this curve (black) and the best-fit vision scaling function, k_v_, from the multimodal dynamic model (green). Dynamic scaling in the vision condition is subtle, and is only cleanly observed close to the stripe on trials following periods of darkness. (F) Mean multisensory condition maximal deviation +/-SEM (purple) plotted as a function of stimulus condition on the previous trial for the 11 flies from Figure 3. Dashed red line is the mean +/-SEM maximal deviation for these same flies on vision trials. The maximal deviation is largest following lights-off (wind) trials, and smaller for both trial types in which the lights were on, suggesting that the vision weight starts low following a period of darkness and increases over time.

Our integration model performs three transformations on incoming sensory information: dynamic scaling, turn command generation, and summation. We first asked if we could infer the relative ordering of these transformations as they take place within the brain - a neural “order of operations” For example, the model described in Figure 5A is mathematically equivalent to dynamic scaling *before* summation, but this need not be the case. We therefore produced a series of multisensory integration models that capture alternative hypotheses about the brain’s information processing hierarchy.

**Table 1:** “Multimodal Dynamic” model best-fit parameters for various stimulus intensities. We used these parameters to generate the data shown in Figures 5-7.

| Simulation Description | Relevant Figs. | $\alpha_w$ (°/s) | $\beta_w$ | $\tau_w$ (s) | $\alpha_v$ (°/s) | $\beta_v$ | $\tau_v$ (s) |
| --- | --- | --- | --- | --- | --- | --- | --- |
| Best-fit (weak vision) | 5; S4 | 50 | 0.15 | 2.3 | 4 | 1.5 | 1.8 |
| Stronger vision, high wind | 6F-H; 7D; S6 | 50 | 0.15 | 2.3 | 8 | 1.5 | 1.8 |
| Stronger vision, medium wind | 6F-H; S5; S6 | 35 | 0.15 | 2.3 | 8 | 1.5 | 1.8 |
| Stronger vision, low wind | 6F-H; S6 | 25 | 0.15 | 2.3 | 8 | 1.5 | 1.8 |

To fit the parameters of these models, we simulated responses to 8 initial orientations, and recovered both wind and visual D-functions from these simulated trajectories (Fig. S4A). In addition, we computed the toward-0° turn rate for simulated multisensory trials (Fig. S4B). Using nonlinear regression, we minimized the error between these simulated measures and those from the empirical data shown in Figures 1 and 2. We found that our three “upstream scaling” models, where dynamic scaling occurs before summation, provided a good fit to these data (Fig. 5B, left). These models differ only in which modality is dynamically scaled. Importantly, each of these models generated a biphasic toward-0° turn rate time course for the multisensory condition, indicating the presence of sequential turning behavior. The root mean squared error of the upwind turn rate time course was similar across all three upstream scaling models (Fig. 5C).

Unlike the upstream scaling models, alternative orders of operation failed to recapitulate our behavioral results. We found that a “time invariant” integration model, with no dynamic scaling, was unable to generate sequential turning behavior in the multisensory condition. The simulated multisensory condition toward-0° turn rate was not biphasic (Fig. 5B, right), meaning that virtual flies only turned downwind on multisensory trials, and did not return to the stripe. This finding confirms our intuition that some component of the core summation model must vary over time to create the sequential turns seen in real flies. To produce a model in which dynamic scaling occurs *after* summation, we added back a dynamic scaling term, but applied it equally to both modalities. This “downstream scaling” model also failed to generate sequential behavior (Fig. 5B,C), indicating that a time-varying component alone is insufficient to recapitulate the empirical data. Critically, the failure of this model tells us that the relative weights of each modality must vary *with respect to one another*. The failure of the downstream scaling model therefore suggests a neural “order of operations, where incoming sensory information across multiple modalities is first scaled, then summed (Fig. 5D).

Our simulations suggest that independent and dynamic scaling of at least one stream of incoming sensory information must occur prior to summation of stimulus-driven turn commands. To determine which upstream scaling model best represented the behavior of real flies, we returned to our empirical data. First, we examined turn rates as a function of time in the wind condition (Fig. 5E, left). We found that the speed of turns through fixed orientations decreased over time in the trial, consistent with the idea that the wind weight decreases over time. The rate of decrease observed experimentally nearly perfectly matched the decay rate obtained by fitting our model. We next sought evidence of dynamic scaling during vision trials. Although we initially found that time in the trial did not influence turn rate, sorting trials by preceding trial type revealed that vision trials preceded by a “lights-off” (wind) trial showed a small increase in turn rate over time, but only for orientations close to the stimulus (Fig 5E, right). Additionally, the sequential response was strongest on multisensory trials preceded by “lights-off” trials (Fig. 5F). The weaker sequential response following “lights-on” trials is consistent with the idea that the vision weight grows slowly after the lights turn on. Based on these analyses, we concluded that the multimodal dynamic model (Fig. 5A,D), where the weights of both modalities change over time, is most consistent with our data, with the two modalities scaled at very different rates and over different magnitudes. The best-fit parameters for this model are shown in Table 1.

To test the validity of our model, we asked if it could recapitulate the measures of multimodal summation and sequential behavior that we observed in real flies (Fig. 3). We simulated flies at 0° and 90° and added a noise term (Fig. 5A; not present for model fitting), which had spectral properties identical to the behavioral data. We found that simulated trajectories closely resembled the empirical data and quantitatively capture our measures of both multimodal summation and sequential behavior (Fig. S5). Our model allows us to draw a number of conclusions about the neural processes underlying the integration of visual and mechanosensory navigational cues (Fig 5D). First, differential dynamic scaling of incoming sensory information is required to produce the turning sequences performed by real flies faced with conflicting cues. Second, this scaling must occur prior to summation so that each modality can be independently re-weighted. Finally, both sensory streams are dynamically scaled, but with different timecourses and magnitudes.

### Multimodal summation and sequential behavior vary continuously with stimulus intensity

Previous studies have found that, contrary to our model, multisensory decision-making often proceeds in “winner-take-all” fashion, with a single stimulus modality dominating behavioral output (Song et al., 2017; Burgos-Robles et al., 2017). In our model, both modalities continuously influence behavior according to their intensity and history. However, it is possible that the dynamic summation we describe occurs only when stimulus intensities are closely matched, and that when one stimulus is much stronger than the other, a winner-take-all strategy emerges. We therefore asked whether our model could generalize to other stimulus intensities. To do this, we presented flies with single and multimodal stimuli at higher and lower windspeeds (*N* = 11, 11, and 12 for wind at 10, 25, and 45 cm/s, respectively), while holding visual stimulus intensity constant.

The trajectories of individual flies suggest that winner-take-all strategies do not emerge even when one stimulus is much stronger than the other. Instead, increasing wind intensity results in stronger signs of multimodal summation and sequential behavior. For example, turns toward the stripe from 90° are fast in the vision condition, but become slower as windspeed increases (Fig. 6A). Similarly, flies fixate near the stripe when started at 0° in the vision condition, but make larger turns away from the stripe as windspeed increases (Fig. 6B). Across flies, correlations emerged when we evaluated turning rate (Fig. 6C; *R*^2^ = 0.08, *p* < 0.05) and latency (Fig. 6D; *R*^2^ = 0.22, *p* < 0.0001) as a function of windspeed for 90° trials, with more intense wind being associated with slower turns at longer latencies. On 0° trials, flies’ maximal deviations over the first 10 seconds of each trial also increased with windspeed (Fig. 6E; *R*^2^ = 0.29, *p* < 0.0001). Considering only the endpoints of each trial might suggest a winner-take-all strategy, but this is an oversimplification. For example, at a windspeed of 45 cm/s, flies end the trial dispersed from 0°, suggesting that wind has won (Fig. 6A,B). However, slow turns back toward the stripe are still visible, implying that the vision command influences turning even at high windspeed. Together, these results argue that dynamic integration of multimodal stimuli during navigation is continuous over stimulus intensity, and that nonlinear winner-take-all dynamics do not apply.

**Figure 6.**
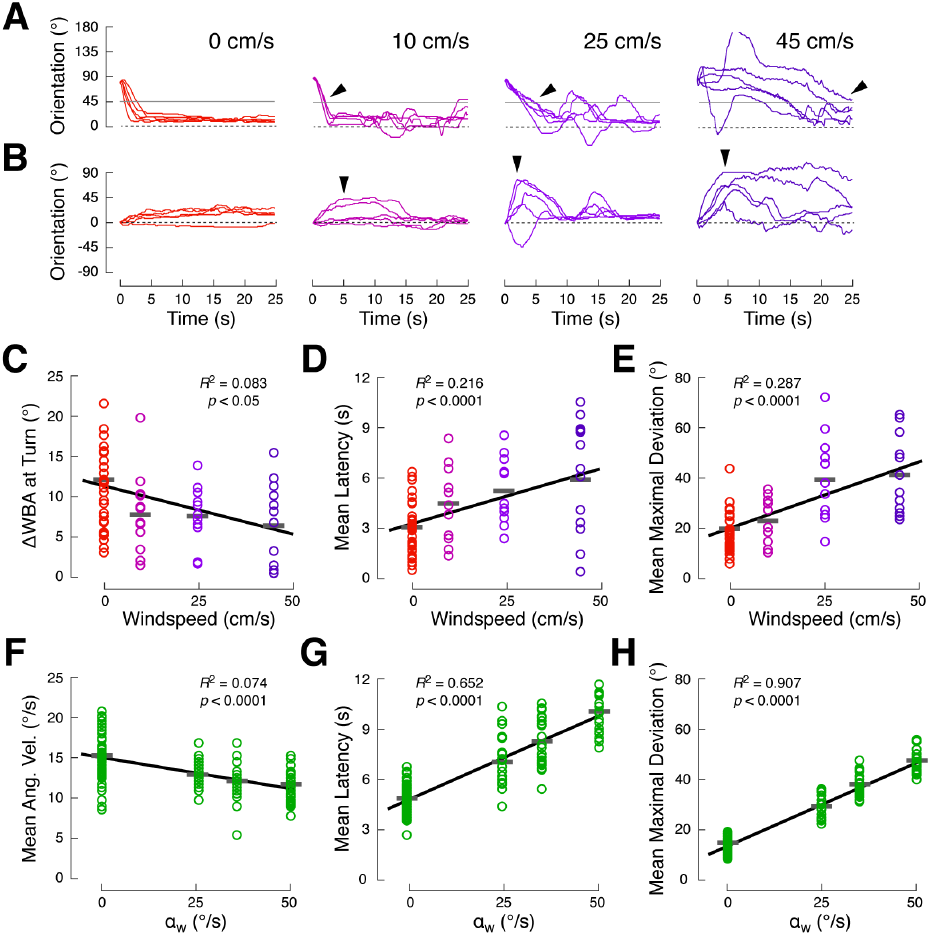
Multimodal summation and sequential behavior vary continuously with stimulus intensity. (A) Slowing of visually-guided turns in the presence of a competing wind stimulus increases with windspeed. Each plot shows 5 trials from single flies beginning at 90°. The leftmost plot represents the vision condition (red), while the right hand plots show the multisensory condition at low (10 cm/s, magenta), medium (25 cm/s, purple), and high (45 cm/s, indigo) wind speeds (each plot is a different experiment/fly). Arrows indicate turns through 45° (gray line). (B) Transient deviations from 0° (arrows) grow with windspeed. Each plot shows 5 trials from single flies beginning at 0°. (C) Mean turn rate through 45° (as in Fig. 3C) as a function of windspeed on 90° trials. Circles: single flies; gray bars: mean across flies. Turn rate is negatively correlated with windspeed (*R*^2^ = 0.08, *p* < 0.05). Black line indicates best linear fit. Each fly was presented with the vision stimulus (*N* = 34) and one of 3 multisensory condition wind speeds (*N* = 11, 11, and 12 for wind at 10, 25, and 45 cm/s, respectively). (D) Latency to turn through 45° (as in Fig. 3C) as a function of windspeed. Visually-guided turn latency is positively correlated with windspeed (*R*^2^ = 0.22, *p* < 0.0001). (E) Mean maximal deviation from 0° (as in Fig. 3G) as a function of windspeed. Flies began each trial at 0° as shown in (B). Maximal deviation is positively correlated with windspeed (*R*^2^ = 0.29, *p* < 0.0001). (F-H) Behavioral measures made on simulated trajectories using the model in Fig. 5A. Each simulated fly was presented with the vision condition (*N* = 36) and one of three multisensory condition wind speeds (*N* = 12 for each). (F) Turn rate (as in (C)) is negatively correlated with windspeed in model simulations (*R*^2^ = 0.07, *p* < 0.0001). (**G**) Latency to turn (as in (D)) is positively correlated with windspeed in model simulations (*R*^2^ = 0.65, *p* < 0.0001). (H) Mean maximal deviation from 0° (as in (E)) is positively correlated with windspeed in model simulations (*R*^2^ = 0.91, *p* < 0.0001).

We next tested the ability of the multimodal dynamic model, which was fit to only a single windspeed, to predict the relationships between stimulus intensity and turn rate, latency, and maximal deviation. To do this, we varied a single model parameter, α_w_, which determines the initial magnitude of the wind D-function, while keeping all other model parameters and relationships fixed. We fit values for α_w_ to the toward-0° turn rate in the wind condition (Fig. S6), then simulated the multisensory condition using these parameters (Table 1). For simulated flies beginning at 90° (*N* = 12 for all α_w_ values), we found the expected relationships between virtual windspeed and turn rate (Fig. 6F; *R*^2^ = 0.07, *p* < 0.0001) and between virtual windspeed and turn latency (Fig. 6G; *R*^2^ = 0.65, *p* < 0.0001), with slower and later visually-guided turns occurring with stronger wind. Similarly, simulated flies beginning at 0° turned farther from the stimulus as virtual windspeed increased (Fig. 6H; *R*^2^ = 0.91, *p* < 0.0001). Due to the simplicity of our model, virtual windspeed is a better predictor of turn rate, turn latency, and maximal deviation in the model than actual wind speed is in real flies. However, the slopes of the best fit lines are quite well matched between the model and real data (Fig 6C vs. F, D vs. G, E vs. H), suggesting that the model makes near-quantitative predictions of the impact that stimulus intensity has on behavior. Overall, these results indicate that, as our model predicts, multimodal summation during navigation occurs continuously over a range of stimulus intensities, and not according to nonlinear winner-take-all dynamics.

### A “sum of turn commands” model can predict responses to both conflicting and synergistic multimodal cues

It is possible that our dynamic summation of turn commands model only applies in cases of multisensory competition. To assess the ability of our model to predict behavior in response to synergistic multimodal cues, we asked whether it could generalize in space. In our stimulus arena, we rotated the black stripe so that it was midway between the upwind and downwind tubes (Fig. 7A). In this arrangement, intuition suggests that a fly approaching the stripe from the upwind side of the arena (making a “downwind” visual turn), will experience synergistic wind and visual turn commands, while a fly approaching from the downwind end will experience conflicting signals. Formally, we can use our model to predict turn velocities by circularly shifting the wind D-function relative to the visual D-function (Fig. 7B), making it clear that downwind-directed turns in the multisensory condition should result from turn commands of the same sign for both modalities, leading to faster turns toward the stripe. In contrast, upwind-directed turns should result from oppositely-signed commands, leading to slower turns. We therefore predicted that flies starting at −90° (facing upwind) should make faster turns toward the stripe at 0° on multisensory versus vision trials, while flies starting at 90° (facing downwind) should make slower turns.

**Figure 7.**
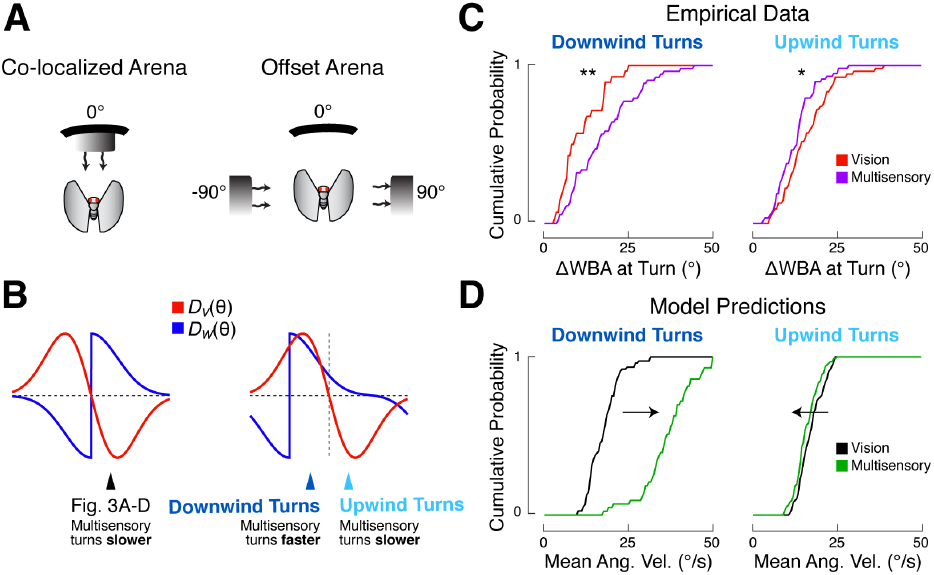
A “sum of turn commands” model can predict responses to both conflicting and synergistic multimodal cues. (A) Schematics of co-localized and offset arena configurations (not to scale). (B) Schematics of the single modality D-functions produced by each configuration. In the co-localized arena, presentation of both stimuli always produces conflicting (oppositely signed) turn commands. In the offset arena, presentation of both stimuli can be synergistic (same signs) or antagonistic (oppositely signed), depending on the orientation of the fly. Downwind multisensory turns are predicted to be synergistic (faster than vision) while upwind multisensory turns are predicted to be antagonistic (slower than vision). (C) Distributions of experimentally measured turn rates in the offset arena for vision (red) and multisensory (purple) trials, sorted by turn direction. Flies were started at either −90° or 90° relative to the stripe to produce downwind or upwind turns, respectively. Rates of individual turns calculated as in Fig. 3C. Downwind turns (left) are faster in the multisensory condition (*n* = 53) than in the vision condition (*n* = 28), as seen in the right-shifted multisensory CDF (rank-sum test, *p* < 0.01). Upwind turns (right) are slower in the multisensory condition (*n* = 59) compared to the vision condition (*n* = 54), resulting in a left-shifted multisensory CDF (rank-sum test, *p* < 0.05). (D) Formal model predictions for the offset arena. Simulation parameters were the same as in Fig. 5E-H, but the wind D-function was circularly shifted by −90° to match the offset arena configuration, as shown in (B). The distribution of turn rates for simulated flies are plotted as CDFs for the vision (black) and multisensory (green) conditions. Rates of individual turns (*n* = 55 for each direction-condition pair) calculated as in Fig. 5G. The distribution of multisensory turn rates is right-shifted compared to vision for the downwind direction (left panel), but left-shifted for the upwind direction (right panel).

Experimentally, we found that flies’ behavior matched these qualitative predictions (Fig. 7C). Behavior in the modified arena was considerably more variable than in previous experiments, perhaps because the tubes that create the wind stimulus act as competing visual targets. As a result, not all flies made the expected turn toward the stripe at 0°. Instead of performing paired comparisons, we therefore evaluated turn rate distributions as independent observations. We collected turn rates for 12 flies navigating in the offset arena, and split the data based on direction of approach to the visual target (“upwind” or “downwind”) and stimulus condition. We then plotted cumulative density functions (CDFs) for all downwind-directed turns toward the stripe in the multisensory (*n* = 53 turns) and vision (*n* = 28 turns) conditions (Fig. 7C, left). Consistent with our predictions, we observed a significant right-shift in the distribution of turn rates on multisensory versus vision trials (*p* < 0.01), indicating that multisensory turns were faster. In contrast, the distribution of upwind-directed multisensory turns (*n* = 59; 54 for vision) was slightly left-shifted, indicating that they were slower (Fig. 7C, right; p < 0.05).

To compare flies’ behavior to the predictions of our model, we used the circularly shifted D-functions to generate synthetic trajectories starting at either −90° or 90°, and analyzed the resulting turns just as for the behavioral data. Similar to our behavioral observations, we found that downwind-directed turns were faster in the multisensory condition than in the visual condition, while upwind-directed turns were slower (Fig. 7D). However, predicted downwind turn velocities were much faster than those we observed behaviorally, likely because there is a ceiling imposed by our device on how fast a turn can be.

Our data from this experiment suggests two conclusions. First, our model can predict the dynamics of turns resulting from both competing and synergistic multimodal cues. Specifically, *conflict* among multisensory turn commands, and not sensation of multiple stimuli alone, is required for the reduction in turn rate that we saw in earlier experiments (Fig 3A-D). Multisensory turns are slowed when the summed turn commands have opposite signs, but are faster if summed commands have the same sign. Second, our proposed computational framework not only generalizes over stimulus intensity, but also over space. We find that a model in which signals from different modalities are first dynamically scaled, then summed as turn commands, can predict the behavior of real flies regardless of how a second stimulus is oriented relative to the first. Together these findings support the idea that a dynamically weighted sum of turn commands can account for many aspects of multisensory integration during navigation.

## Discussion

### Summation of dynamically weighted turn commands underlies multisensory orienting

In this study we used a combination of behavioral monitoring and simulations to identify computations underlying multisensory integration during navigation. We found that flies faced with competing visual and mechanosensory cues exhibited two behavioral phenomena. First, flies turned toward a visual target more slowly in the presence of a competing wind stimulus. Second, flies’ responses to competing stimuli were dynamic, with early-trial turns dominated by wind and late-trial orientation dominated by vision. We were able to marry these disparate responses under a single computational framework in which incoming sensory streams are first dynamically scaled in time, translated into position-dependent turn commands, and summed (Fig. 5D). Our model could recapitulate many aspects of the observed behavior and was robust to changes in stimulus intensity and spatial arrangement.

Based on the mathematical structure of our model, we drew a number of conclusions about the neural processing hierarchy that underlies multisensory orienting. First, we used the failure of alternative models to argue that a dynamic scaling step must exist, and must be upstream of summation so that the relative weights of stimuli can change over time. Dynamic scaling could result from neural processes both before and after turn command generation, but we can conclude that the scaling process must be completed prior to summation. We also used the structure of our model to argue that summation takes place on turn commands, and not on some other signal, such as the sensory signals themselves, turn probabilities, or the target orientations prescribed by each stimulus. Direct summation of sensory signals is not consistent with our data, as we found that wind could make vision-guided turns faster or slower depending on the orientation of the fly relative to the wind source (Fig. 7). Summation of turn probabilities, a model recently advanced by another group (Gepner et al. 2015), is also unlikely, as we found that not only turn probability, but also turn rate, depend on the position and intensity of the stimuli (Figs. 6 and 7). It seems plausible that the disparity between these studies reflects the differences between two forms of navigation, one comprised of oriented turns toward spatially localized stimuli, and the other based on stochastic re-orientations in response to spatially diffuse odor and light. Finally, weighted summation of multimodal target orientations, a hypothesis proposed in previous work (Budick, Reiser and Dickinson, 2007), is also disfavored by our data. Such a model, which does not specify how an animal should proceed from its current orientation to the target, requires a separate motor planning step downstream of summation. If the motor plan is constructed downstream of cue integration, then the speed of turning should depend only on the location of the target orientation relative to the fly, not on the locations of the individual sensory stimuli. Our observation that flies proceed from 90° to a target orientation of 0° at different speeds in the presence or absence of wind (Fig. 3) is inconsistent with this model. Based on these considerations, we conclude that integration most likely takes place on turn commands, in units of angular velocity.

Our model builds on a classical account of stripe fixation in tethered flies (Reichardt and Poggio, 1976), which found that closed-loop orientation to visual stimuli could be predicted from a combination of position- and motion-based D-functions measured in open-loop. Similarly, we find that multimodal closed-loop orientation can be predicted from open-loop D-functions (measured at the beginning of trials), combined with scaling terms fit to the data. In our model, we have not included a visual or mechanosensory motion term. This may account for the relatively poor ability of the model to fit the detailed dynamics of responses in the vision condition (Figs. S4 & S5). Other models of *Drosophila* optomotor behavior rely exclusively on stimulus velocity (Reiser and Dickinson, 2013; Mongeau and Frye, 2017). However, certain classes of visually-driven turns in flies do seem to depend on integrated stimulus position, and not stimulus velocity (Reichardt and Poggio, 1976; Bender and Dickinson, 2006; Schnell et al., 2014). Addition of a motion-based term may improve the model fit to our vision data, but at the cost of additional complexity.

### Advantages of a dynamic sum of turn commands model

An advantage of a navigation model in which multimodal cues are integrated in units of angular velocity is that it can predict both turn dynamics *and* the behavioral end-points that were previously described as a weighted sum of target orientations. To ask whether our model can generate intermediate target orientations in response to conflicting cues, we simulated responses to spatially offset wind and visual stimuli, but omitted the dynamic scaling process that causes vision to dominate toward the end of trials (Fig. S7). We found that our model could indeed produce intermediate steady-state orientations that resemble a linear combination of wind intensity, visual intensity, and stimulus position, consistent with previous work (Müller and Wehner, 2007; Budick, Reiser and Dickinson, 2007). This observation suggests that summation of turn commands could represent a general strategy for integrating cues from multiple sensory modalities during orienting behavior.

At the neural level, summation of angular velocities represents a simpler strategy than summation of target orientations. Neural D-functions could be constructed from spatially tuned sensory neurons that activate premotor populations to drive directed turns. For example, to build an attractive visual D-function, neurons selective for ipsilateral stimuli could excite neurons that produce ipsilateral turns. Downstream pooling of this premotor activity could produce multisensory summation without direct interaction of sensory signals. Conversely, the specification and subsequent summation of real-world target orientations would require an additional unit transformation, from egocentric to allocentric, before summation could occur.

Because it can be constructed from canonical neural computations, our model suggests obvious targets for investigating the neural underpinnings of multisensory orienting. Recent studies have identified a likely locus for the integration of multisensory turn commands in the *Drosophila* brain. The central complex, a region of the insect brain implicated in navigation behavior (Strauss and Heisenberg, 1993; Ofstad, Zuker and Reiser, 2011; Kahsai, Martin and Winther, 2010; Seelig and Jayaraman, 2015), is known to receive both ipsilaterally-tuned input from visual interneurons (Omoto et al., 2017; Sun et al., 2017) and directional mechanosensory signals (Ritzmann, Ridgel and Pollack, 2008; Phillips-Portillo, 2012). Neurons in several regions of the central complex have been implicated in turning behavior (Green et al., 2017; Turner-Evans et al., 2017; Martin et al., 2015), suggesting that this structure might translate directional sensory stimuli into oriented movements. The laminar organization of the central complex (Hanesch et al. 1989; Wolff et al., 2015) could provide an anatomical substrate for turn commands derived from different modalities to be aligned and summed, similar to the alignment of sensory signals that drive gaze or head orientation changes in the vertebrate tectum (Lee, Rohrer and Sparks, 1988; Carello and Krazulis, 2004). For these reasons, the central complex represents fertile ground for future tests of our multimodal summation model.

### Asymmetric filtering of sensory input as a mechanism for behavioral sequence generation

We found that flies faced with competing cues reliably executed a turning sequence, responding first to the aversive wind stimulus, then to the attractive visual target. Sequential behavior has been previously observed in navigating insects, and is particularly prevalent in mosquito host-finding behavior. Carbon dioxide, detected in plumes over long distances, gates and drives heat-seeking behavior (McMeniman et al., 2014), host odor attraction (Lacey, Ray and Cardé, 2014), and attraction to discrete visual objects (van Breugel et al., 2015). This navigational sequence — CO_2_ tracking, then visual object attraction, then heat- and host odor-seeking — arises primarily due to the sequence of cues encountered by the navigating insect, not on any direct neural coupling between these sensorimotor units (van Breugel et al., 2015).

In contrast, the sequential response we observed must arise from sequential activity generated in the brain. In our paradigm, light turns on slightly faster than wind (Fig. 1B), so the early wind turns we observed under multisensory conditions cannot arise from the sequence of stimulus encounters. We also observed that the behavioral sequence is abolished when the antenna are stabilized, arguing that it arises only when neural signals from both modalities are present. Importantly, we know that the antennae faithfully encode wind without physically adapting (Yorozu et al., 2009), arguing against the possibility that dynamic scaling and sequential responses arise from antennal biomechanics. Together, these observations support a neural origin for the observed turning sequence.

The neural and computational frameworks that can produce serial activation of motor motifs have traditionally been thought to lie in one of two categories: feedforward excitatory chains, as seen in zebra finch song and human speech production (Long et al., 2010; Houghton and Hartley, 1995), or hierarchical suppression (Davis, 1979) of premotor motifs that are driven in parallel (Averbeck et al., 2002). Recent work in *Drosophila* has also identified lateral and feedback disinhibitory circuit motifs that support behavioral sequence transitions and promote sequence integrity (Jovanic et al., 2016). Asymmetries in sensory gain have been proposed as a possible model for sequence generation *in silico* (Seeds et al., 2014) but have not, to our knowledge, been demonstrated experimentally. Our results provide evidence that early, asymmetric processing of incoming sensory data can indeed produce sequential behavior. Our work therefore contributes to the view that a wide range of computations and neural circuit motifs are capable of generating simple behavioral sequences.

Why might flies want to execute a sequence in response to conflicting wind and visual stimuli? The two stimuli we used for this study have different ethological meanings to a fly. Distant visual landmarks are often used by insects to stabilize heading (Graham and Collett, 2002; Reiser and Dickinson, 2010) and may be used to generate extended segments of straight flight. In contrast, the type of rapid mechanosensory stimulus we present here would most likely arise from a gust of wind, and would in natural flight be accompanied by changes in optic flow as the fly is displaced by the wind. In the absence of that optic flow, the mechanosensory stimulus may be discounted over time. Dynamically decreasing the weight given to this stimulus would allow the fly to ignore the distracting wind and focus on its visually-driven navigational goals. Consistent with this interpretation, we found that presentation of longer, dynamic wind stimuli resulted in a gradual decrease in the response to wind onset (data not shown).

### Stimulus-guided navigation as a behavioral window into the neural computations that support multisensory integration

The question of how the nervous system integrates, represents, and utilizes sensory data from multiple modalities has interested neuroscientists for decades. A major focus of many previous studies has been to generate quantitative models of how information from different modalities is integrated over time to drive behavioral decisions (Meredith and Stein, 1983; Raposo et al. 2014). However, because the decision itself is localized in time, the dynamical processes occurring in the brain must be inferred. A complementary approach is to study a behavioral output that is itself a continuous function of time. This approach takes advantage of the many natural behaviors, such as foraging, locomotion, and migration, that evolve continuously in time, and permits direct observation of behavioral dynamics. The ability to quantitatively describe how stimuli are translated into ongoing behavior has proven to be a fruitful approach for making direct links between neural mechanisms and behavior (Gray, Hill and Bargmann, 2005; Coen et al. 2014; Gepner et al. 2015; Shulze et al. 2015).

In this study we used orientation behavior of flying Drosophila to study the integration of conflicting or synergistic mechanosensory and visual stimuli. We found that behavior could be described as a continuous sum of dynamically weighted, stimulus-driven turn signals. Importantly, we found that the relative weight given to a modality varies with that modality’s intensity (Fig. 6). Due to these weighting properties, our model closely resembles others that have been proposed to describe integration during multisensory decision-making (Ernst and Banks, 2002; Fetsch et al., 2012; Drugowitsch et al. 2014). Specifically, those studies suggest that cues from different modalities are summed weighted by their reliability, for which intensity is a proxy. These results raise the intriguing possibility that the computations underlying orientation behavior in a simple genetically-tractable model organism may be closely related to the computations performed by more complex organisms during multisensory decision-making.

## Acknowledgements

We would like to thank Karla Kaun and Matthieu Louis for flies and Marc Gershow, Michael Long, Michael Reiser, and Dmitry Rinberg for feedback on the manuscript. Members of the Nagel and Schoppik labs provided additional feedback and helpful discussion. This work was supported by grants from NIH (R00DC012065), NSF (IOS-1555933), the Klingenstien-Simons Foundation, the Sloan Foundation, and the McKnight Foundation to K.I.N.

### Author Contributions

Conceptualization, T.A.C. and K.I.N.; Methodology, T.A.C. and K.I.N.; Software, T.A.C.; Investigation, T.A.C.; Formal Analysis, T.A.C.; Visualization, T.A.C.; Writing – Original Draft, T.A.C.; Writing – Review & Editing, T.A.C. and K.I.N.; Funding Acquisition, K.I.N.; Supervision, K.I.N.

### Declaration of Interests

The authors declare no competing interests.

## Materials & Methods

### Flies

Unless otherwise specified, all flies were raised at 25°C on a cornmeal-agar-based medium under a 12 hour light/dark cycle. All experiments were performed on adult female flies 3-5 days post-eclosion. The data from the first round of experiments (Figs. 1 & 2) described in this paper came from control flies in a genetic silencing screen. These flies were raised at 18°C and were of the following genotypes:

w- CS; UAS-TNTe/+; R12D12-gal4/tubp-gal80^ts^

w- CS; UAS-TNTe/+; R14C07-gal4/tubp-gal80^ts^

w- CS; UAS-TNTe/+; R24C07-gal4/tubp-gal80^ts^

w- CS; UAS-TNTe/+; R24E05-gal4/tubp-gal80^ts^

w- CS; +; R24E05-gal4/tubp-gal80^ts^

w- CS; UAS-TNTe/+; R28D01-gal4/tubp-gal80^ts^

w- CS; UAS-TNTe/+; R29A11-gal4/tubp-gal80^ts^

w- CS; UAS-TNTe/+; R34F06-gal4/tubp-gal80^ts^

w- CS; UAS-TNTe/+; R38B06-gal4/tubp-gal80^ts^

w- CS; UAS-TNTe/+; R38H02-gal4/tubp-gal80^ts^

w- CS; UAS-TNTe/+; R43D09-gal4/tubp-gal80^ts^

w- CS; UAS-TNTe/+; R46G06-gal4/tubp-gal80^ts^

w- CS; UAS-TNTe/+; R60D05-gal4/tubp-gal80^ts^

w- CS; +; R60D05-gal4/tubp-gal80^ts^

w- CS; UAS-TNTe/+; R65C03-gal4/tubp-gal80^ts^

w- CS; +; R65C03-gal4/tubp-gal80^ts^

w- CS; UAS-TNTe/+; R67B06-gal4/tubp-gal80^ts^

w- CS; UAS-TNTe/+; R73A06-gal4/tubp-gal80^ts^

w- CS; +; R73A06-gal4/tubp-gal80^ts^

w- CS; UAS-TNTe/+; R84C10-gal4/tubp-gal80^ts^

w- CS; +; R84C10-gal4/tubp-gal80^ts^

w- CS; UAS-TNTe/+; tubp-gal80^ts^/+

Because these flies were raised at 18°C, expression of gal4 was restricted by gal80. We therefore treated these flies as near-wild-type controls for the purposes of this paper, and pooled data from all genotypes. In all other experiments, flies were Canton-S (CS) wild-type. Note that all findings in non-CS flies were replicated in CS wild type animals.

All flies were cold anesthetized on ice for approximately 5 minutes during tethering. A drop of UV-cured glue (KOA 30, Kemxert) was used to tether the notum of anesthetized flies to the end of a tungsten pin (A-M Systems, # 716000). Tethered flies’ heads were therefore free to move. For antennal stabilization experiments, a small drop of the same UV-cured glue was then placed between the second and third segments of the fly’s antennae, as depicted in Figure 4A. Flies were then allowed to recover from anesthesia for 30-45 minutes in a humidified chamber at 25°C before behavioral testing. All experiments were performed at 25°C.

### Behavioral apparatus

The behavioral apparatus and the software that controlled it were both custom made. The stimulus arena consisted of a circular piece of rigid plastic (1.25” dia, 1.25” high) lined on the inside with white copy paper. On opposing faces of the arena, holes were drilled 0.5” from the top of the arena to accommodate two teflon tubes (ID 3/16”, OD 1/4”, Cole-Parmer, # 06605-32) that delivered the wind stimulus. The “upwind” tube was packed with smaller 18 gauge stainless steel tubes (Monoject, # 8881-250016) to laminarize flow at the fly. For all experiments except the final one, a 0.3” wide strip of black tape (Pro-Gaff) was vertically oriented behind the upwind tube, as shown in Figure 1A. For the final experiment (Fig. 7), the strip of tape was positioned 90° from the upwind tube. Both tubes extended 1/2” into the arena, leaving a 1/4” gap where the fly would sit. A 3D-printed tube holder stabilized the tubing and the arena to maintain a rigid position during rotation. Six IR LEDs (870 nm, Vishay, # TSFF5210) provided illumination for wing imaging. They were arranged in a circular pattern on the floor of the arena and were each aimed toward the fly.

A combination rotary union / electrical slip ring system (Dynamic Sealing Technologies, # LT-2141-OF-ES12-F1-F2-C2) converted stationary electrical and pressurized air inputs outside the arena into rotating electrical and pressurized air connections inside it, facilitating stimulus delivery. The rotary union was coupled to the stimulus arena and tube holder with a flanged adapter (NEMA23 mount, Servocity). Tygon tubing of various diameters (ID 1/16” - 5/16”, Fisher) sequentially carried house air through a regulator (Cole-Parmer, # MIR2NA), a charcoal filter (Dri-Rite, VWR, # 26800), a mass flow controller (Aalborg, # GFC17), a three-way solenoid valve (Lee, # LHDA1233115HA), and the rotary union before air was delivered to the fly. The downwind tube was connected to house vacuum through the rotary union, a second identical solenoid valve and a flowmeter (Cole-Parmer, # PMK1-010608). Flow of house air and vacuum were always switched on and off together.

Each tethered fly was held in place with a pin holder (Siskiyou) which was lowered into the stimulus arena. Tethered flies were positioned directly between the upwind and downwind tubes, approximately 0.1” from the wind source, and were held at a pitch angle of approximately 30 degrees with respect to the horizon, near their natural flight posture. Behavioral imaging was performed by a camera (Allied, # GPF 031B) and VZM 100i zoom lens lens (Edmund Optics, # 59-805) equipped with an IR filter (Edmund Optics, # BP850-22.5) that was positioned 1.5 ft above the fly. Strobing of the IR LEDs below the fly was coordinated with the camera frame rate (50 Hz) via a microcontroller (TeensyDuino, PJRC.com), ensuring that IR illumination did not heat the fly over the course of an experiment. Images of behaving flies were digitally processed by a custom LabVIEW (National Instruments) script to produce a nearly binarized white fly on black background. This image was used to extract the angle of each fly’s wings with respect to its long body axis. The difference between wing angles was multiplied by a static gain (0.04) and converted into an angular velocity on each sample. This gain was selected through a set of pilot experiments, where this setting was found to allow flies to fixate a stripe without orientation “ringing.” A static offset was manually selected at the beginning of each experiment to correct for subtle yaw rotations (less than 5°) relative to the camera. The difference between wing angles over time was saved for later analysis. All communication between LabVIEW and arena components was mediated by a NIDAQ board (National Instruments, # PCIe-6321).

The angular velocity signal calculated in LabVIEW was sent to a stepper motor controller (Oriental, # CVD524-K) that converted this information into stepping commands. These commands drove a stepper motor (Oriental, # PKP566FMN24A) by an appropriate amount on each imaging sample. The motor stepped at 500 Hz, and was capable of driving rotations of the arena up to 162 °/s. We used the built-in smooth drive and command filter functionality of the motor controller to smooth the drive and reduce vibrations. The motor drive was transferred to the stimulus arena with a pair of gears (Servocity, # 615238), closing the loop back to the fly. Although rapid body saccades could not be reproduced by this motor/gain combination (Muijres et al., 2015), slower course adjustments are well captured. We were restricted in possible rotational velocities by the competing demands of reasonable behavior, minimal vibration, and the high torque required to rotate the pressurized rotary union. The behavioral gain and maximal rotational velocity chosen represent a balance between each of these considerations.

### Stimulus design

The visual stimulus, as described above, occupied approximately 30° of visual angle. We controlled ambient illumination of the stripe by placing the entire rig in a foam-core (ULINE, # S-12859) box that was then draped with a black curtain (Fabric.com), rendering the arena light-tight. A set of warm white LED strips (HitLights, # LS3528_WW600, 36 SMDs) along the roof of the box were powered by a combination driver/ dimmer (Luxdrive 2100mA Buckblock, LED Dynamics) and controlled via a microcontroller. Illumination of the stripe was measured with a power meter (Thor, # PM100D). Light levels used in these experiments were 0.1, 0.25, and 15 μW/cm^2^, measured at a wavelength of 488 nm.

The wind stimulus, as described, was manipulated via a mass flow controller and a solenoid valve. Charcoal-filtered house air was pushed out of the upwind tube while vacuum was pulled through the downwind tube. Our custom LabView script regulated the flow through the MFC, and the valves were used to create rapid onset/offset kinetics (see Fig. 1). Flow at the fly, regulated by the MFC, was calibrated with an anemometer (a Dantec MiniCTA with split fiber film probe, # 55R55) that had previously been calibrated under similar conditions. Windspeeds of 0, 10, 25 and 45 cm/s were used in our experiments. We also used the anemometer to verify that wind intensity was constant across orientations of the stimulus arena as it rotated (see Fig. 1). After initial calibrations, we re-checked the wind stimulus every 3-4 months to ensure stimulus consistency over time.

### Experimental design

All experiments were controlled via custom LabVIEW scripts running on a desktop computer. In the experiments shown in Figs. 1 & 2, each fly underwent 32 trials that were each a pseudorandomized combination of stimulus condition and initial orientation. There were four stimulus conditions (wind only, light only, both together, or no stimulus) and eight starting orientations (one every 45°). For these first experiments, wind speed was set to 45 cm/s and light intensity was set to either 0.1 or 0.25 μW/cm^2^. At the start of each trial, the stepper motor rotated the arena to the initial orientation selected by the LabView script, and the selected stimulus/stimuli were switched on. The fly was then given 25 seconds of control over the orientation of the arena. At the end of each trial, all stimuli were switched off, and the arena ceased to rotate. The next set of starting conditions were then selected, and the subsequent trial would begin. The inter-trial interval (ITI) was dictated primarily by the amount of time it took the motor to position the arena at the next starting orientation. The ITIs were therefore variable, but all fell within the range of 50-500 ms.

In the experiments shown in Figs. S2 and S3, the sequence of events that comprised each trial were identical to the first experiment, but the four stimulus conditions were changed. For this experiment, our LabView script selected among four wind speeds (0, 10, 25 or 45 cm/s), meaning three possible stimulus selections were wind-only conditions, and one possible stimulus selection was a no-stimulus condition. There were still 8 starting positions, and therefore still 32 total trials. A command signal from the LabVIEW script to the MFC altered the windspeed during the ITI, and the solenoid valve was opened at the start of the trial. This allowed us to vary wind intensity while maintaining rapid onset/offset kinetics.

In the experiments shown in Figs. 3 and 4, we retained the trial design but altered both the initial orientations set and the stimulus conditions set. Here we used only two initial orientations (0° or 90°) and three stimulus conditions (wind only, light only, or both together), for a total of 6 possible stimuli. Each stimulus was forced-selected 5 times per fly, but the order of stimulus presentation was random. A full experimental session was therefore comprised of 30 trials. In this experiment, we used a windspeed of 25 cm/s and an ambient light intensity of 15 μW/cm^2^.

The experiments shown in Fig. 6 were identical to the those in Figs. 3 and 4, but with altered windspeeds. Any given fly experienced wind of either 10 or 45 cm/s in the wind-only and multisensory conditions. Light intensity was unchanged across the third experiment (15 μW/cm^2^).

The experiment shown in Fig. 7 was identical to the third experiment with the exception that the stimulus arena was spatially rearranged and the initial orientations were altered to compensate for this re-arrangement. The white copy paper and black tape that comprised the high-contrast visual stimulus (see above) were replaced such that the black bar was now oriented 90° away from the wind source (see Fig. 8a). The stripe remained at 0°, the upwind tube was positioned at −90°, and the downwind tube at 90°. The possible initial orientations used were −90° or 90°. Wind intensity was 45 cm/s, and ambient illumination at the fly was 15 μW/cm^2^.

### Analysis of behavioral data

All analyses were performed using custom Matlab (Mathworks, Natick, MA) scripts. All analyses were performed on one of two metrics: the raw difference between wing angles, as converted to angular velocity by multiplying by a static gain (*g*, 0.4, Eq. 1), and the online-integrated orientation of the fly relative to the stimulus.

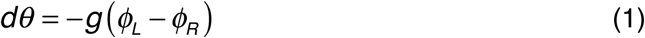

Unless otherwise noted, all measures derived from the difference in wing angles used a filtered version (Low-pass Butterworth filter, 2 Hz cutoff frequency). All correlations were tested for significance using a permutation test on Pearson’s *R*, all comparisons between stimulus conditions in single flies used paired *t*-tests and all comparisons across flies used the Wilcoxon rank-sum test, unless otherwise indicated.

D-functions for each stimulus were calculated by splitting all trials for each fly by starting orientation, then taking each fly’s average angular velocity over the first two seconds of each trial. Data was then pooled across flies and plotted. The two second duration was chosen to maximize the differences between orientations: using a longer or shorter window does not change the general shape of the D-functions, but does change their magnitudes. For the wind and multisensory D-functions, the data was split for the 0° starting condition by taking two averages across flies: one average for all mean-positive angular velocities, and one for all mean-negative data points. This allows us to highlight the aversive nature of the responses, and is necessary because flies can turn either direction to make downwind progress. If we do not split the data at 0°, the population mean is very close to 0 °/s, which implies that head-on wind does not induce flies to turn. This is not the case, as flies tend to turn *fastest* when presented with head-on wind. To show this, we plotted for each stimulus condition the mean turn rate of each fly for early turns starting at 90° as a function of mean turn rate starting at 0° (Fig. S1). When flies fell far below the diagonal, indicating faster turns from 0° than from 90°, we split the D-function at 0° for that stimulus condition. The visual D-function was not split in this manner as flies turn faster from 90° than from 0° (most data falls above the diagonal, see Fig S1). Finally, D-function “peaks” (see Figs. S2 and S3) were the angular velocities corresponding to the 0° starting position for each fly.

Orientation histograms were calculated as the mean normalized occupancy across flies for each of 72 heading bins (each 5° wide, with the first bin centered on 0°) over the last 15 seconds of each trial. We split trials by stimulus condition, but not by initial orientation. We normalized these distributions and treated them as probability density functions. The last 15 seconds of the trial were chosen because more than 95% of flies had completed their stimulus onset-driven turn by this time.

Similarly, quadrant preference measures were also calculated as the mean normalized occupancy across flies for each of 4 heading bins (90° wide, with the first bin centered on 0°) over the last 15 seconds of each trial, split by stimulus condition. We called the heading bin centered on 0° “upwind,” the one centered on 180° “downwind,” and the other two bins “crosswind”. The null hypothesis for these preferences is therefore 1:1:2 for upwind:downwind:crosswind.

We developed the toward-0° turn rate metric to evaluate whether a fly is turning toward the stripe and/ or upwind tube, which were both located at 0°. Such turns are positive under this measure, and turns away from 0° are negative. To calculate toward-0° turn rate, we applied the following formula (Eq. 2, unsimplified for algorithmic clarity) to the raw difference in wingbeat angles for each time step on each fly.

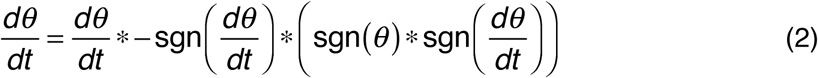

We first ask if the fly is moving away from 0° by taking the product of the sign of orientation and the sign of angular velocity: this returns a 1 if the signs are the same, indicating a turn away from 0°. We then ask if angular velocity is negative by taking the negative sign of angular velocity. We then take the product of these two “checks,” which yields 1 or -1, and multiply angular velocity by this value. Functionally, this formula flips the sign of turns toward 0° from 0° < θ< 180° and turns away from 0° from 0° >θ > -180° while leaving the signs of other turns unchanged. This means that all turns toward 0° are now positive, and all turns away from 0° are now negative. We low-pass filtered (0.5 Hz cutoff) the resulting time courses.

Analysis of the 0°-90° protocol (Figs. 3-6) included a few additional measures. For trials in which flies began at 90°, we asked whether they crossed a threshold half-way to the visual or wind targets (0° or 180°, respectively). Thus, the visual threshold was at 45°, and the wind threshold at 135°. For trials in which flies did cross a threshold, we calculated the local turn rate at the threshold as the mean angular velocity in a 1 sec window centered on the threshold crossing. The timing of the threshold crossing became our latency measure. For each fly, we then took an average across trials of the local turn rate and latency measures. The number of trials that contributed to this average varied for each fly, though we discarded flies that completed fewer than 3 threshold crossings to prevent single trials from being over-weighted in the population means. The threshold crossing discussed throughout this paper for the multisensory condition is the visual threshold of 45°.

For trials in which flies began at 0°, we found the maximal orientation reached in the first 10 seconds of each trial. This window was chosen to capture the early turning that characterized the sequential response, while ignoring late-trial turns that sometimes occurred. We took the mean of this maximal orientation for each fly in each condition. The individual fly means therefore always included 5 trials for this measure.

Analysis of the spatially offset stimulus protocol (Fig. 7) began by defining visual turns as either upwind or downwind in direction. For consistency with previous experiments (Figs. 4-6), we retained the 45° orientation threshold for visually-guided turn definition. Because the visual target was positioned at 0° and the wind at −90°, an upwind visual turn was defined as the mean local (+/-0.5 s) angular velocity of a fly with a negative angular velocity while crossing 45°. Conversely, a downwind visual turn was defined as the mean local angular velocity of a fly with a positive angular velocity while crossing -45°. This measure captured the same turns as the paired analysis described for the 0°-90° protocol (see above), but allows for individual flies to not make the expected turn. This measure also considers all eligible turns that meet these definitions, instead of only the first turn, as in earlier analyses (Figs. 3-6). This analysis yielded an upwind and downwind turn distribution for each stimulus condition that we plot as CDFs in Figure 7.

### Model of multisensory orienting

We developed a computational model to ask whether the diverse behaviors we observed experimentally could be understood in terms of a single simple framework. The core of our model is a sum of single modality position-dependent turn commands (D-functions), inspired by the observation that addition of a wind stimulus changes the rate of turning towards the visual stripe. To generate sequential behavior, we allowed the strength of each D function to vary over time. The full version of the model includes dynamic scaling terms for each modality (“multimodal dynamic”). We also tested simpler versions in which only one modality varies over time (“dynamic wind” and “dynamic vision”), both vary together with the same time course (“downstream scaling”), or neither varies (“time invariant”).

In the multimodal dynamic model, angular velocity is determined by Equation 3 (Fig. 5A).

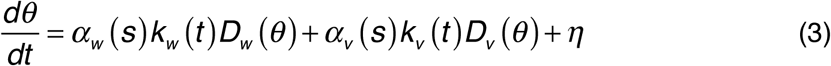

The two D-functions, D_v_(θ) and D_w_(θ) were unitless and had idealized shapes motivated by our experimental data (Fig. 1). We modeled the wind D-function, D_w_(θ) as the normalized product of a step function from -1 to 1 at 0° and a Gaussian with a standard deviation of 50° (Eq. 4). We modeled the visual D-function, D_v_(θ) as the normalized product of a line with a slope of 1 s^-1^ and a Gaussian with a standard deviation of 50°(Eq. 5).

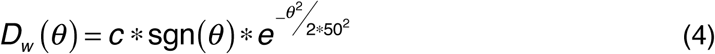

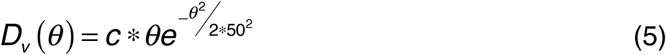

Note that *c* for each equation is simply the normalizing constant equal to one over the max value of the rest of the function evaluated between 0° and 180°. The normalizing constant has units of °^-1^ to make the D-functions unitless. Both of these idealized D-functions are plotted in Figure 5A.

The functions k_s_(t) are unitless dynamic scaling terms that modify the strength of each modality’s turning drive over time. We took these functions to be exponentials (Eq. 6), increasing for vision and decaying for wind.

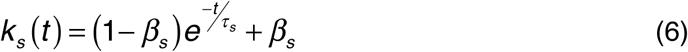

These scaling functions are structured to start with a value of 1 at *t* = 0 and end at a fixed fraction, parameterized by β_s_, at *t* = ∞. If β_s_ is larger than 1, the influence of that stimulus will grow over time, and if β_s_less than one, it will decay. To achieve a growing visual influence and a decaying wind influence, we restricted β_v_ to be larger than 1 and β_w_ to be less than 1 during subsequent fitting.

The magnitude of each D-function was given by α_s_(s), a static function that increases non-linearly with stimulus intensity, and is zero when the associated stimulus is not present (Fig. S5). α_s_(s) has units of °/s. Thus, each modality contributes a turn command that depends on stimulus intensity, the orientation of the stimulus with respect to the fly, and the time since stimulus onset. The turn commands for each modality sum. For simulated trials (but not for model fitting) we also added a noise term (ƞ, see Eq. 3). We generated the noise for each trial by multiplying white noise in the frequency domain by the power spectrum of the empirical no stimulus condition angular velocity signal, then returning this product to the time domain. This procedure gave us a noise time course on each trial that matched the spectral properties of empirical data in the no stimulus condition.

To simulate responses of the model, we generated trials organized identically to trials in our experiments, with three or four conditions and two or eight starting orientations, depending on the experiment, per simulated “fly.”The size of the time step in the simulation was matched to the behavioral sampling rate (50 Hz, dt = 0.02 sec). At each time step we used the current orientation of the fly to calculate the angular velocity, then updated the orientation for the next time step. To simulate stimulus conditions, we simply modified α_s_(s) terms as appropriate. In the wind condition, α_w_ had a nonzero value and αv was set to 0. Conversely, α_w_ was set to 0 and α_v_ had a nonzero value in the vision condition. Both α_w_ and α_v_ were nonzero in the multisensory condition.

Free parameters of the multimodal dynamic model are the three terms, α, βand τ(Eqs. 3 & 6) for each modality. To fit these parameters, we ran deterministic simulations (lacking the noise term), and computed the resulting single modality D-functions and a down-sampled version of the multisensory condition toward-0° turn rate, just as we did for our experimental data. We used nonlinear regression to minimize the mean squared error between the model results and the empirical data shown in Figures 1 and 2. Using the best fit parameters (Table 1), we then ran a full simulation of 100 virtual flies in the non-deterministic full model. The results of this full simulation are plotted against the empirical data in Figure S4 (right column). We refer to non-deterministic simulations run using the parameters obtained by fitting to a deterministic model as “best-fit simulations”.

As described above, and as shown in Figure 1, the visual stimulus intensity in the first experiment was low compared to the strength of the wind. In order to model the 0°-90° paradigm and generate the plots in Figure 5E-H, we therefore needed an increased visual weight (α_v_), just as we increased visual stimulus intensity in real flies. We did this by fitting model results to the vision condition toward-0° turn rate data using the same procedure described above (Fig S5; Table 1), resulting in a larger α_v_. All other parameters were kept at their initial best-fit values. We analyzed trajectories from this “stronger vision” parameter set identically to the empirical data, pulling out the same metrics of turning rate at threshold, turn latency, and maximal deviations.

The equations describing the simpler versions of the model are shown below for the time invariant (Eq. 7) and downstream scaling (Eqs. 8 and 9) models. In the time invariant model (Eq. 7), no dynamic scaling functions (k_s_) are used.

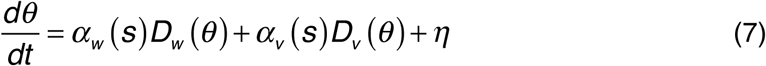

In the downstream scaling model (Eqs. 8 and 9), βd may take any value from 0 to ∞; there are no restrictions on whether overall stimulus influence should grow or decay with time.

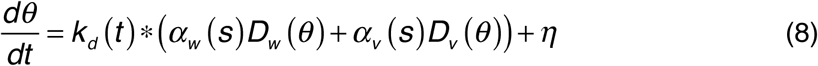

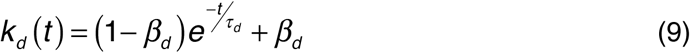

The single modality dynamic scaling models are shown below in Equation 10. In the dynamic vision model (Eq. 10, top), only k_v_ is preserved from the multimodal dynamic model, while in the dynamic wind model (Eq. 10, bottom), only k_w_ is preserved.

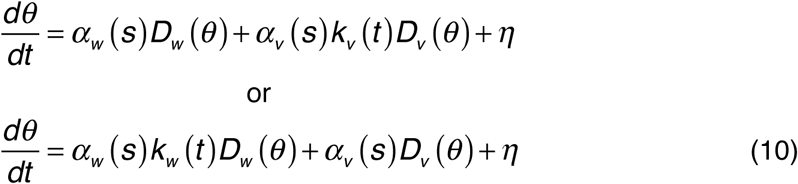

We used the same fitting procedure to find the best fit parameters for these models to the data from experiment 1. We calculated the root mean squared error between the D-functions and multisensory condition upwind turn rate data from each best-fit simulation and those from the empirical data, allowing us to evaluate the performance of each model relative to the others (Fig. S4).

To differentiate between the various upstream scaling models, we asked for each modality whether we could see evidence for dynamic scaling in the absolute turn rate through a given orientation as a function of time. By examining a constant orientation throughout the trial, we could directly probe if flies modulate their turn velocity over time. We pooled data from all flies (*N* = 120) in the first experiment (Figs. 1 and 2), and used 12 equally-sampled time bins (same number of turns per bin) to evaluate mean turn rate through the chosen orientation in each bin. For the wind condition, we evaluated a range of orientations (30°, 45°, 60°, 75°, 90°, 105°, 120°, 135°, and 150°). For the vision condition, we examined turn rate through 45°, the threshold we defined for visual turn latency in previous analyses, on trials preceded by wind trials (when the lights were off). The change in visual scale is subtle, and can only be seen close to the stripe following periods of darkness, while the change in wind scale is robust and can be seen generally. We fit the absolute turn rate time courses obtained in this manner with exponential curves, and evaluated the parameters of these exponentials in relation to the best-fit *k* functions in the multimodal dynamic model.

For the varied windspeed simulations, we retained the “stronger vision” α_v_ parameter because we used the same higher light intensity in the actual experiments (see Fig. 6). We found a best-fit α_w_ for each intensity by fitting the simulated wind condition toward-0° turn rate to the empirical data using the fitting procedure described above (Fig. S5; Table 1). All other parameters were fixed across windspeeds.

Finally, we simulated the effect of rotating one stimulus 90° relative to the other by circularly shifting the wind D-function while keeping the vision D-function unchanged (Fig. 7B). We used the “high wind” parameter set (Table 1) to match the relevant stimulus intensities, and analyzed visually-guided virtual turns as described above for the stimulus offset experiments.

